# Phages associated with horses provide new insights into the dominance of lateral gene transfer in virulent bacteriophages evolution in natural systems

**DOI:** 10.1101/542787

**Authors:** V.V. Babenko, A.K. Golomidova, P.A. Ivanov, M.A. Letarova, E.E. Kulikov, A.I. Manolov, N.S. Prokhorov, E.S. Kostrukova, D.M. Matyushkina, A.G. Prilipov, S. Maslov, I.S. Belalov, M.R.J.C. Clokie, A.V. Letarov

## Abstract

Tailed bacteriophages (Caudovirales order) are omnipresent on our planet. Their impressive ecological and evolutionary success largely relies on the bacteriophage potential to adapt to great variety of the environmental conditions found in the Biosphere. It is believed that the adaptation of bacteriophages, including short time scale adaptation, is achieved almost exclusively *via* the (micro)evolution processes. In order to analyze the major mechanisms driving adaptation of phage genomes in a natural habitat we used comparative genomics of G7C-like coliphage isolates obtained during 7 years period from the feces of the horses belonging to a local population. The data suggest that even at this relatively short time scale the impact of various recombination events overwhelms the impact of the accumulation of point mutations. The access to the large pool of the genes of a complex microbial and viral community of the animal gut had major effect on the evolutionary trajectories of these phages. Thus the “real world” bacteriophage evolution mechanisms may differ significantly from those observed in the simplified laboratory model systems.

## Introduction

The tailed bacteriophages are the most numerous, diverse and widely spread group of viruses known (Clokie et al. 2011). Their ecological success suggests that they are particularly good at adapting to a wide range of environmental conditions. It is generally accepted that bacteriophage adaptation to the environment occurs by microevolution (or short-term evolution). The evolution of bacteriophage genomes has generally been studied by comparative genetics and more recently on comparative genomics of natural phage isolates (Marston and Martiny 2016; Ha and Denver 2018; Sazinas et al. 2018). These efforts revealed that phage evolution is driven by combination of the accumulation of mutations and by the modular exchange of genetic material with the external sources such as other phages or the host cell genome (Mavrich and Hatfull 2017; Maddamsetti et al. 2018; Meier-Kolthoff et al. 2018; Petrie et al. 2018)

Early studies performed on lambdoid temperate phages (Botstein 1980) gave a rise to the concept of phage genomes as the mosaic constructs build of distinct genetic modules. This theory was further developed to suggest that all phage genomes in the biosphere can access a common pool whereby genes and modules can be swapped (Hendrix et al. 1999). After more sequence data became available, the concept of the core genome and an accessory genome emerged (Comeau et al. 2007; Chan et al. 2014). In this premise, core-genome regions are thought to evolve by the accumulation of point mutations rather than by lateral modular transfer. This is augmented by a peripheral genome where lateral transfer events are frequently observed. Large scale pairwise comparisons of sequenced phage genomes led Mavrich and Hatfull (Mavrich and Hatfull 2017) to the conclusion that phages fall into two distinct lifestyle groups where one has low and one has high lateral gene content flux (i.e. with predominant vertical or modular evolution).

Phage adaptation *via* co-evolution with bacterial hosts has also been extensively studied laboratory settings whereas relatively stable co-existence of phage and bacterial populations during multiple passages is generally observed (Scanlan et al. 2011; Morley et al. 2017; Scanlan et al. 2017), see also reviews (Buckling and Brockhurst 2012; Golais et al. 2013). Although the details of the dynamics of co-evolutionary changes of phage and bacterial phenotypes vary depending on experimental design, in all the cases where the genetic background of phage adaptation was analysed, it was found that the phages generally evolved by accumulating point mutations or, less frequently, small insertions and deletions in genes that encode for their receptor recognition proteins (Meyer et al. 2012; Akusobi et al. 2018; Petrie et al. 2018), see also reviews (Scanlan et al. 2011; Letarov and Kulikov 2017).

The relative simplicity of laboratory-based experiments however could actually be responsible for driving these observed types of changes. This is because in artificial settings, the genetic pool to which phages can access is limited to the genome of the host strain and to highly related genomes of phage lineages that are formed *in situ.* The relative impacts of mutation accumulation and lateral gene transfer during short-term microevolutionary adaptation of the phage genomes in natural habitats are still poorly understood and in this study, we aimed to redress this by examining an alternative study system. We performed a comparative genomic study of seven field isolates of coliphages highly related to the bacteriophage G7C (Kulikov et al. 2012) that were obtained from horse feces from horses stabled in the same geographical location. The idea behind this approach is that if we can observe this ‘rare’ phage type from within in a closed system then we can determine the evolutionary pressures in a non-lab limited, ‘real world’ evolutionary context. The period of the phage isolation spans 7 years and therefore the microevolutionary dynamics of this group of phages can be observed in a distinct ecological system (i.e. the local horse population) over this time period.

The choice of horse intestinal microbiome as an experimental system for this study has multiple advantages to study natural phage evolution. The surface to volume ratio in horse intestines is greater than in small animals such as mice and this limits the influence of the macro-organism can impose over the microbial community. In addition, the typical 48-72h food retention time in horses is much longer then typical time gaps between food intake and defecation acts, so the horse intestinal ecosystem acts as a natural, stable chemostat. Importantly the stability and complexity of *E. coli* populations is very useful and convenient for follow up culture-based work. This stability is due to the large volume of the caecum and lower intestine being populated by metabolically active microbes that digest cellulose (Julliand and Grimm 2016). This contrasts with smaller animals where the active part of the microbiome is associated with mucosa (Poulsen et al. 1995).

Interestingly and in contrast to other gut systems, almost all the coliphages isolated from horse feces were found to be virulent and genetically diverse, even within individual animals (Golomidova et al. 2007; Kulikov et al. 2014; Golomidova et al. 2016).

The original phages under study - G7C and G8C - were isolated from a single horse faeces sample collected during a study of horse intestinal coliphage ecology study in 2006 (Golomidova et al. 2007; Kulikov et al. 2012). These phages were isolated using the environmental *E. coli* strain 4s a host, which has O22 type of O-antigen with additional glycosylation (Knirel et al. 2015).

G7C and G8C are related to the well-studied bacteriophage N4 (Kazmierczak and Rothman-Denes 2006; Choi et al. 2008) which has a particular organisation of the adsorption locus (Kulikov et al. 2012) consisting of two tail spike proteins gp66 and gp63.1. The function, and structure of these proteins were previously characterized by us (Prokhorov et al. 2017) and gp63.1 was have a rare enzymatic activity O-antigen acetyl esterase, which probably explains why they display a narrow host range. (Kulikov et al. 2007; Golomidova et al. 2007). This further confirms our strategy to screen *E. coli* 4s as a good host on which to identify further G7C-related phages.

Using this strategy, we isolated a further five G7C phages. The comparison of the genomes of these viruses coupled with experimental evaluation of biological effects of some of the observed lateral gene transfer events lead us to conclude that in this ecological system lateral genetic transfer is the dominant mode of evolution. This is in striking contrast to the *in vitro* situation where the co-evolution relies on a very limited number of the selected point mutations. Moreover, the core-genomes of these phages suggests a high level of mosaicism due to the frequent recombination events observed between closely related genomes.

## Results

### Bacteriophage isolation strategy

Five new phages were isolated in this project; Alt63 from an existing lysate and four from new samples from the same population of horses that are focus of this study (from the Equestrian centre in Neskuchny Sad, Moscow, Russia).

Alt63 was isolated from a primary lysate from which we then had previously purified phage G7C (Golomidova et al. 2007). In order to separate it from the dominant G7C phage we grew it on a mutant strain of *E.coli* 4sI, that lacks the O-antigen O-acetylation (Knirel et al. 2015) and is thus. This strain is resistant to phage G7C but so we hypothesized that it would be a useful tool to isolate additional phages.

In order to search for additional G7C-like phages from our horse population we plated horse feces samples on our original host *E. coli* 4s and screened for this phage group using a) PCR to target the virion RNA polymerase gene g50 and b) RFLP analysis of genomic phage DNA. This strategy allowed us to isolate two new phages, St10y and 10Gb in 2010; and a further two phages N4G2 and Sz33 in 2013. Together with Alt63, G7C and G8C these seven phages are termed the G7C-set.

Of particular note to this study the N4 like phages were a feature found within our horse population but not elsewhere, so no N4-like viruses that infect *E.coli* 4s were found either in other locations or from other horse populations tested. This held true for both 2010 or 2013 despite multiple attempts to isolate them.

### The determination and overall comparison of the genomic sequences

The genomes from the six new N4-like phages were fully sequenced and a single contig for each phage was assembled. As all the phages were closely related to G7C, we set the left end of the genome at the conserved nucleotide sequence (TGgGGGGCT) present in this site (Kulikov et al. 2012). As the capsid morphogenetic proteins of all these new phages are almost identical to their G7C homologs, we assumed that the size of the encapsulated DNA in these phages is the same that was found for G7C. Thus, the right end position, and terminal redundancy size was directly calculated for each of these viruses, the sequences deposited in GenBank reflect this.

The nucleotide sequences of G7C-set phages are highly related to each other (Fig. 1A) with the overall level of nucleotide identity >90%. The whole genome phylogeny was reconstructed using an alignment-independent algorithm by JSpecies software analysis (Drummond and Rambaut 2007). JSpecies phylogeny is consistent with the time of the bacteriophage isolation (Fig. 1B). The G7C-set phages clustered together, and are closer to each other than EC1.UPM and PhAPEC7, their closest relative found in GenBank.

**Fig. 1.**
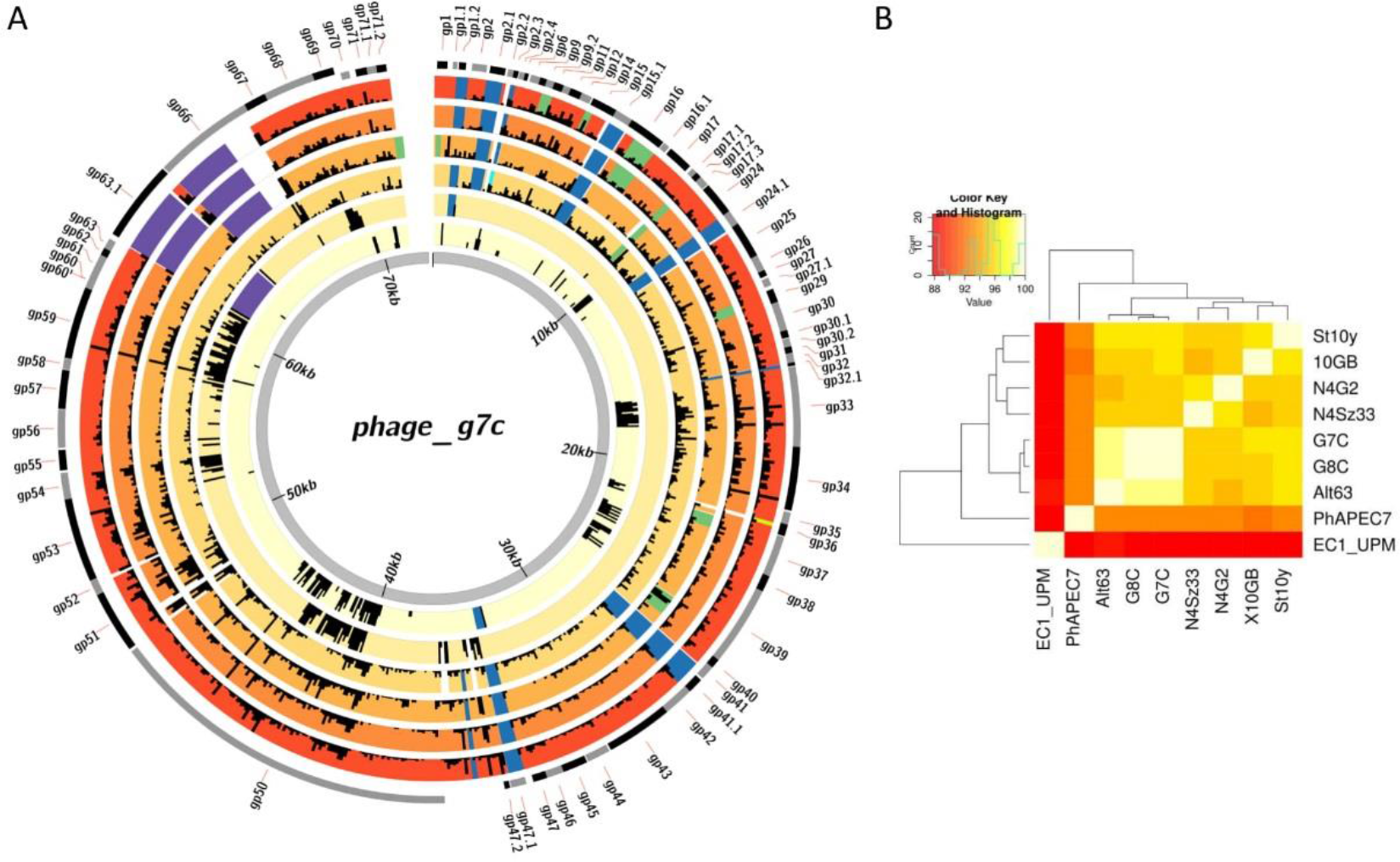
**A.** The diagrams of the SNP density for the alignments of all the phages against G7C genome used as a reference. The background color corresponds to the overall distance of the genomic sequence from G7C phage as calculated by JSpecies software. The window size is 100 b.p. The maximum pin height corresponds to 8 SNPs per window. White areas to non-aligned (containing gaps or the fragments replaced by non-orthologous sequences, except for the C-terminal modules of the tail-spike proteins that are shown in violet). The blue fragments indicate the deletions of the genetic material compared to the G7C sequence, the green fragments indicate the insertions. The coordinates of G7C phage are used. **B.** The phylogenetic tree reconstructed using the whole genome sequences by JSpecies software. Phage EC1.UPM and PhAPEC7 are taken as an outgroup.

### The recombination events

To detect recombination events, we aligned the genome sequences using MAUVE (Darling et al. 2004). The interruptions of sequence homology were detected and manually inspected using the pairwise alignment generated by Martinez-NW algorithm within Lasergene 6.0 software. We limited our search for events that affected more than 100 bp regions, and resulted in less than 50% of the local nucleotide identity with respect to one or more of the phages. Within the G7C-set, we detected 22 loci where genetic material was inserted, deleted, or replaced by non-homologous sequences at least in one of the phages (Table S1 and Fig. 1A).

The majority of the module swapping occurred within ORFs that are not part of the core genome within N4-like phages (Chan et al. 2014). These genes generally either have no assigned function, or are related to homing endonucleases. The exception to this was the genes that encoded for DNA and RNA polymerases, and for tailspikes (Table S1, Fig. 1A).

### Insertions in essential RNA and DNA polymerases genes

As most of the genes where recombination has occurred have no known function, it is difficult to predict the impact of such events. However, it was intriguing that the integration of external genetic material occurred within the RNA-polymerase gene 16 in phages N4G2 and Sz33 and within DNA-polymerase gene 39 in phage 10Gb because these proteins are essential for viability. Thus, further effort was put into establishing how these genes and thus the phages remained functional despite these seemingly detrimental alterations.

In N4G2 and Sz33 ~800 bp insertions interrupted the gene of RNA polymerase 2 subunit A (g16). These insertions are identical and are referred to as the g16 insertion. In phage 10Gb a similar sized but distinct insertion was found in the DNA polymerase gene g39. In both cases, the insertions encode multiple stop-codons that preclude further read through of the genes in any reading frame.

Both these g16 and g39 insertions contain short internal ORFs that are predicted to be the homing nucleases, that are frequently present within introns. In order to analyse if these insertions represent actual introns, we performed the Rfam search (Kalvari et al. 2018) yielded no significant hits the analysis of putative secondary structures of their RNA revealed a that they form a complex pattern of the stem-loop structures. These structures suggest that they may be type I introns (Fig. 2A, B). It is noteworthy that the ends of the insertion are positioned closely in the structures, and the G-residue is located immediately after the predicted splicing point – this is a key characteristic feature of the bacterial type I introns (Hausner et al. 2014).

**Fig. 2.**
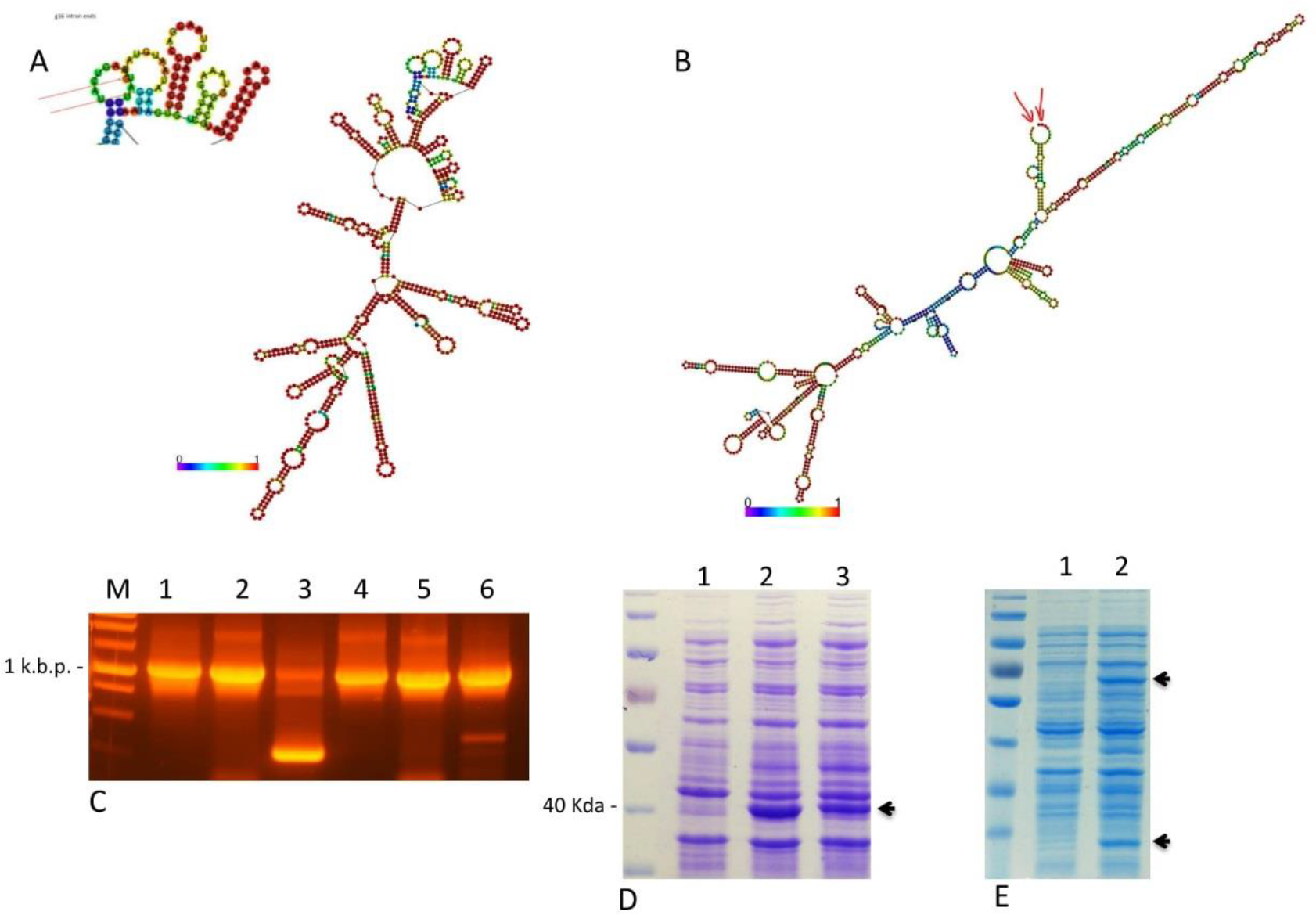
**A.** Predicted secondary structure of the RNA transcribed from 800 b.p g16 and g39 **B.** insertions. Arrows indicate the position of the insertions ends. **C.** Reverse-transcription (RT) PCR analysis of the splicing of g16 оf Sz33 phage and g39 of 10 GB insertions. Lanes: 1 – phage Sz33 DNA, 2-RNA of Sz33 infected cells without RT, 3 – RNA from Sz33 infected cells with RT; 4 phage 10GB DNA, 5-RNA from 10GB infected cells without RT, 6 - RNA from 10GB infected cells with RT. The weak low molecular weight band seen in g39 RT-PCR is due to non-specific amplification of a cellular RNA that was confirmed by the sequencing. **D.** Expression of recombinant gp16 from the intron containing genes. Lanes: 1-empty vector, 2= pN4G2_g16, 3- pSz33_g16. **E.** Expression of recombinant gp39 from the phage 10GB gene containing non-splicable insertion sequence. Lanes: 1 - empty vector, 2-p10GB_39.

To investigate the splicing activity of the intron-like elements we isolated the RNA from the cells infected by the phages Sz33 and 10Gb and performed RT-PCR analysis for both putative introns (Fig. 2C). The splicing of g16 intron was reliably detected, and the splicing point confirmed by the sequencing thus confirming that g16 insertion is an intron.

We also cloned g16 from both N4G2 and SZ33, and expressed the proteins. The size of the recombinant proteins corresponded to the spliced RNA (Fig. 2D) and the peptide containing the transition point (QAVMTAFYGSEAK; the transition point is between a.a. residues shown in bold) was confirmed by MALDI-TOF analysis (data not shown).

In contrast, no traces of spliced product were observed for the g39 insertion in phage 10Gb (Fig. 2C). We therefore conducted studies to determine how this gene is translated. We were initially unable to clone the gene either when intact or when the G7C insertion -free version was used as a PCR template. It thus appeared that this G7C-like DNA polymerase is toxic for *E. coli*. We generated a C-truncated version of gp39 and after multiple attempts; we obtained clones of phage 10Gb g39 in pGEM-T vector under inducible T7 promoter to generate a pG39_10GB plasmid.

The protein expression from this plasmid gave rise to two separate recombinant proteins (Fig. 2E) whose molecular weights corresponded to the N- and C-terminal parts of the gp39 encoded upstream and downstream of the putative intron. This was confirmed by MALDI-TOF analysis (data not shown). Manual inspection of the intron-like sequence revealed the presence of a Shine-Dalgarno sequence the correct distance from the ATG codon that it is likely to be the initiation site of the C-terminal fragment translation.

The re-sequencing of the G39 present in the cloned C-truncated gene on our pG39-10GB plasmid indicated that in there are two mutations in the putative intron sequence (one nucleotide substitution and 1 b.p. deletion breaking the frame of the putative homing nuclease gene). This observation suggests that the insertion sequence in g39 may also feature the toxicity (presumably due to the nuclease activity) and the mutations probably abolished this toxicity.

In summary, the data suggest that the invasion of the non-spliceable mobile sequence into phage 10Gb gene 39 has lead to conversion of N- and C-terminal fragments of the DNA polymerase gp39 into separate subunits. It is not clear if these proteins interact physically to form two-subunit holoenzyme, however perfect viability of the phage indicates that the DNA polymerase is functional *in vivo*.

### The swapped C-terminal domains of the tail spike proteins

It is not surprising that the tailspike proteins were altered by recombination events because proteins involved in host recognition are common targets of modular swapping. Consistent with this, within the G7C-set isolates we observed two allelic variants of gp63.1 and four alleles of gp66 (Fig. 3A). Both these proteins are the tail spikes (Kulikov et al., 2014; Prokhorov et al., 2017). The aa sequences of the N-terminal parts of these different allelic variants are conserved. In contrast, the C-terminal parts of both gp63.1 and gp66 from different allelic types are unrelated, or very distantly related to each other despite the surrounding sequences being highly homologous even at the nucleotide level. The BlastP analysis of aa sequences of the C-terminal parts of all the allelic variants of gp63.1 and gp66 identify multiple phage related proteins for each of them (Kulikov et al. 2014; Prokhorov et al. 2017).

**Fig. 3.**
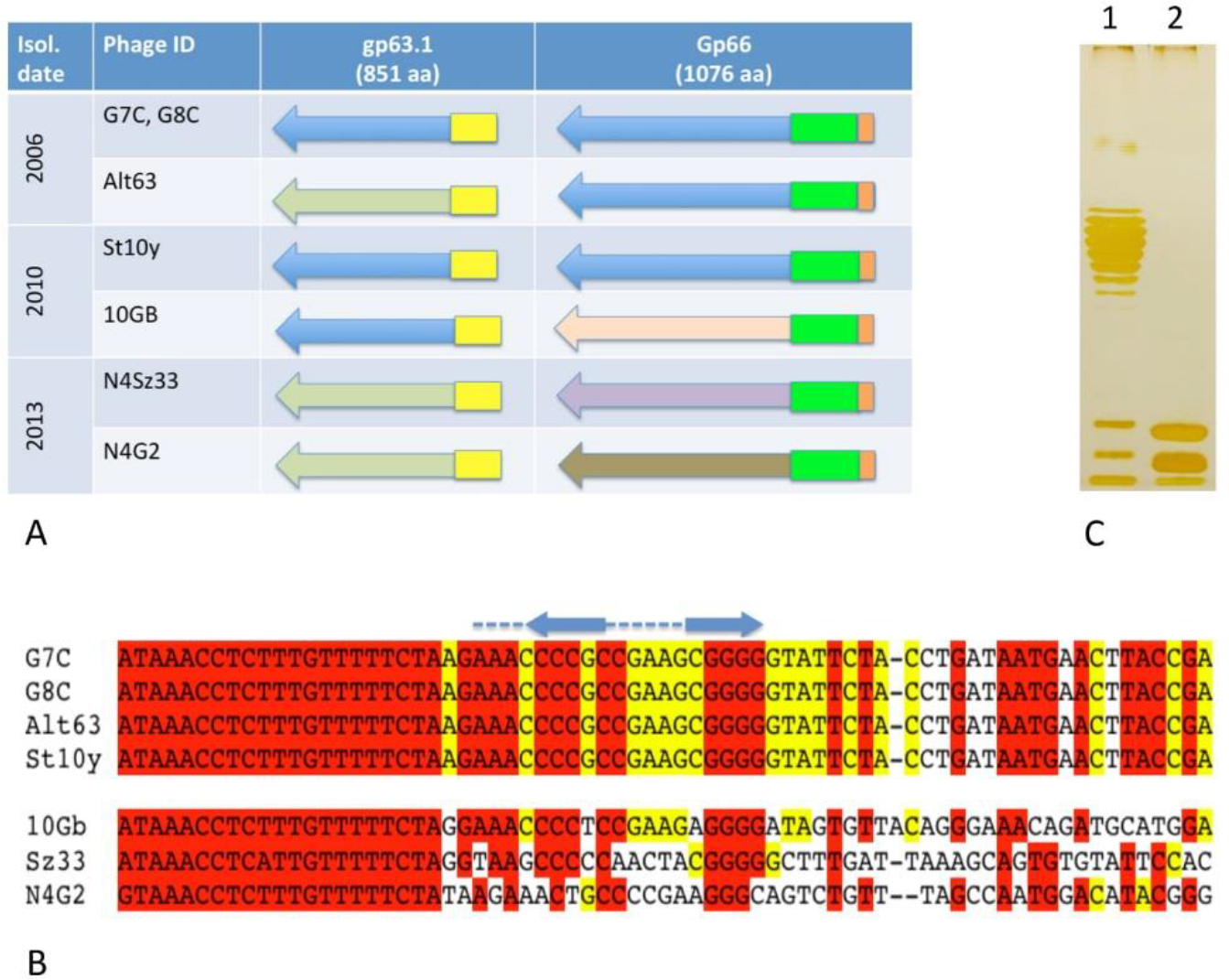
**A.** Allelic variants of gp63.1 and gp66 in G7C-set bacteriophages. The N-terminal parts of the proteins are conserved while the C-terminal moieties are subjected to the modular swapping. **B.** The alignment of the nucleotide sequences adjacent to the left end of g66 including the transcription terminator – like stem loop structure (shown by the blue arrows and lines above) located between genes 63.1 and 66. The phages in the top part contain G7C-like version of gp66, those in the bottom all have different allelic variants of this protein. **C.** The SDS-PAAG analysis of E. coli 4s LPS before (lane 1) and after (lane 2) the treatment by purified gp63.1A protein. The degradation of OPS chain is observed.

In terms of identifying where this source of genetic information has come from, in all cases, the best matches for the loci variants are bacterial proteins presumably encoded by the prophages. The matches were to genomes from *E. coli, Salmonella, Shigella*, and *Enterobacter*. These results suggest that prophages play a significant role in the evolutionary dynamics of the host recognition proteins in G7C-like bacteriophages.

It is not clear if the high frequency of lateral transfer of the genetic information into tail spike genes from unrelated sequences can be explained by intensive selection or there is a mechanism stimulating recombination in this locus. Interestingly, in observed recombination exchanges in g66 were limited at the left hand side (with respect to the standard genomic sequence orientation). This region forms a putative stem-loop structure similar to transcription terminator that was previously identified in G7C genome between genes 63.1 and 66 (Kulikov et al. 2012). This structure appears to be conserved in all the other phages of our set, although in N4G2 this stem-loop sequence appears to be less evident than in the other six phages (Fig. 3B). These observations suggest that this structure may play a specific role in the recombination mechanism mediating the frequent modular exchanges in this point of the genome. It may be a recognition site of some meganuclease(s) mediating homing-like mobility of the genetic modules encoding gp66 C-terminal fragment variants. Alternatively the stem-loop structure may limit the Holiday junction migration and stimulate it processing by the nucleases.

### Gp63.1 protein from phage Alt63 has an OPS depolymerising activity

In order to establish the significance of domain swapping we examined the enzymatic activity associated with the Alt63-type receptor recognition module of gp63.1 tail spike.

All the G7C-set isolated on *E. coli* 4s strain appear to have very narrow host range, which is most likely to be because their infection depends on the specific recognition of the O22-like O-antigen (Knirel et al. 2015; Prokhorov et al. 2017). Indeed, our efforts to find alternative natural hosts for the G7C-set phages among the horse-derived *E. coli* isolates were unsuccessful. However, phage Alt63 was isolated using a *wclK* mutant of *E. coli* 4s that has no O-acetylation of the O-antigen (Knirel et al. 2015), and thus is completely resistant to the G7C phage. Interestingly although we isolated this new phage Alt63 that targets strain 4sI, no phage mutants of the original G7C could be selected for that were able to overcome this resistance (Knirel et al. 2015) These observations indicated indirectly that gp63.1 protein from Alt63 may possess a different type of the enzymatic activity. Phages N4G2 and Sz33 that carry Alt63-type of gp63.1 tailspike are also able to grow on *E. coli* 4sI while G8C, 10Gb and St10y having G7C-type of this protein (deacetylase) grow exclusively on *E. coli 4s* wt as do the prototype of this series phage G7C. So we hypothised that the enlarged host range of Alt63 phage may be linked to the changed enzymatic activity of the tailspike gp63.1. To check this hypothesis we cloned gene 63.1 from Alt63 protein into expression vector with C-terminal His-tag and produced the recombinant protein that we named gp63.1A. The protein was purified using the metal-affinity chromatography. The lipopolysaccharides (LPS) extracted from *E. coli* 4s was incubated with purified protein and analysed by SDS-PAAG electrophoresis (Fig. 3C). The pool of the molecules with variable length of O-polysaccharide (OPS) of variable length was converted to single O-unit – containing molecule population indicating the OPS-depolymerising activity of gp63.1A.

### Detection of hidden mosaicism of the conserved parts of the genome

The modular replacement events described above lead to changes of the genetic content of the affected loci due to deletion of the genetic material or acquisition of unrelated sequences. However, most of the genes are present in all the G7C-set phages thus forming a local core-genome. In order to determine if this core genome evolves mainly vertically due to point mutation accumulation or it features hidden mosaicism streaming out the recombination between highly similar sequences, we performed comparative analyse of the individual phylogenies of the core genes.

To do so we selected all the genes that are present in all seven phages of our collection. In total, 67 sets containing 7 homologous genes each were formed. The sequences were aligned within each of such sets and 67 matrixes of the similarity were generated. The Pearson correlation coefficients between these matrixes were calculated and the similarity matrix was built and visualized as a tree and as a heatmap (Fig. 4). This diagram reflecting the similarity of the clusterization patterns of different genes in our phages (or in simpler words similarity of the topologies of the phylogenetic trees that can be built for each gene) allowed us to define three groups of the genes of the local core-genome.

**Fig 4.**
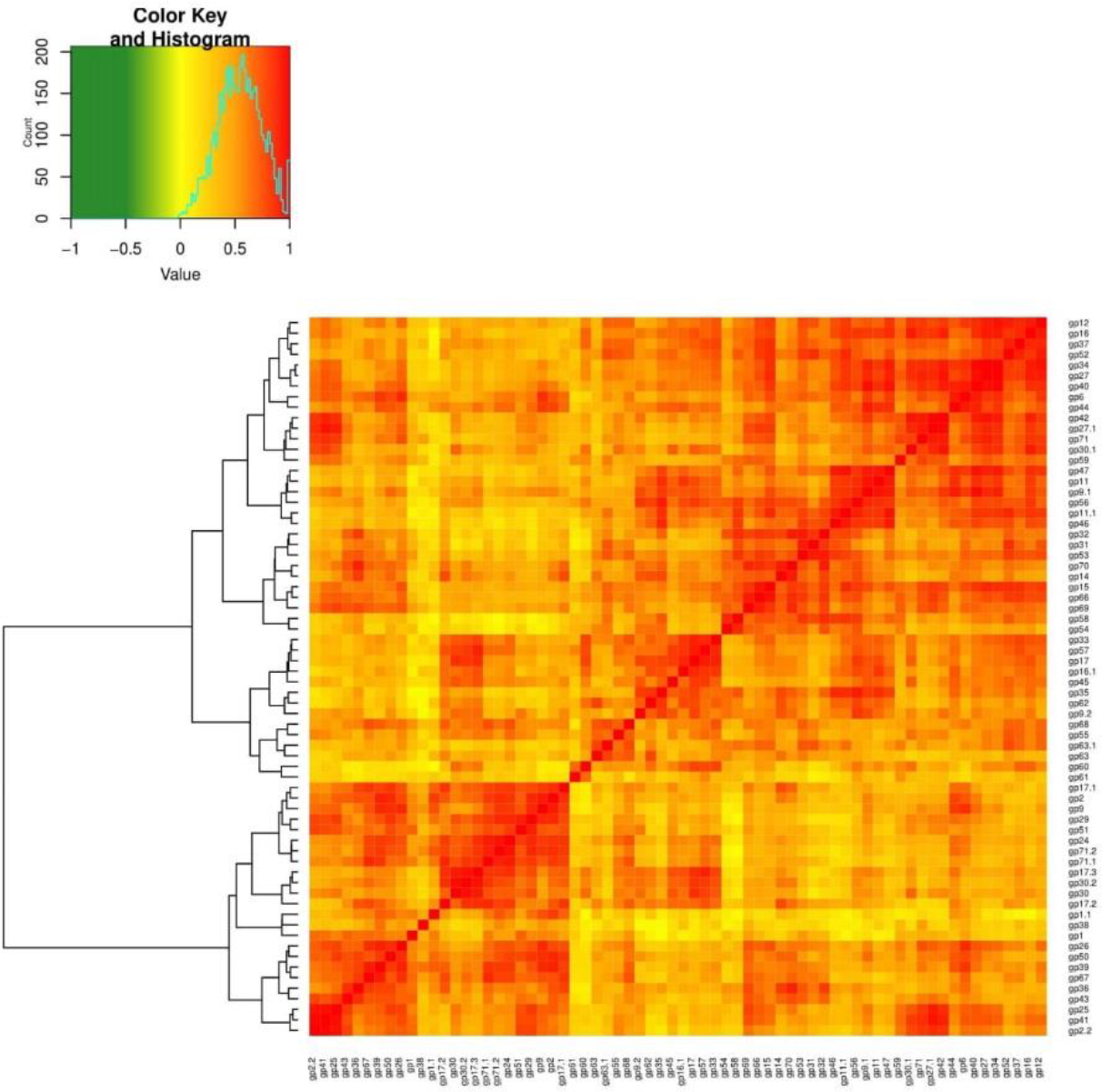
The heatmap of Spearman correlations of the matrixes of similarity built for 67 genes of the local core-genome of G7C-set. Three groups of the local core-genome genes differing by the patterns of similarity (topology of their phylogenetic trees) were defined.

The genes belonging to each group were then marked on the genetic map of the phage G7C (fig. 5A). The distribution of the genes belonging to three identified groups appears mosaic; there is only weak association of these groups with the predicted functions. Group #1 includes the majority of the structural proteins, group #2 is enriched by the nucleic acid metabolism enzymes, while group #3 is miscellaneous of proteins with different predicted functions (replication, host defence evasion, cell lysis) and hypothetical proteins.

**Fig. 5.**
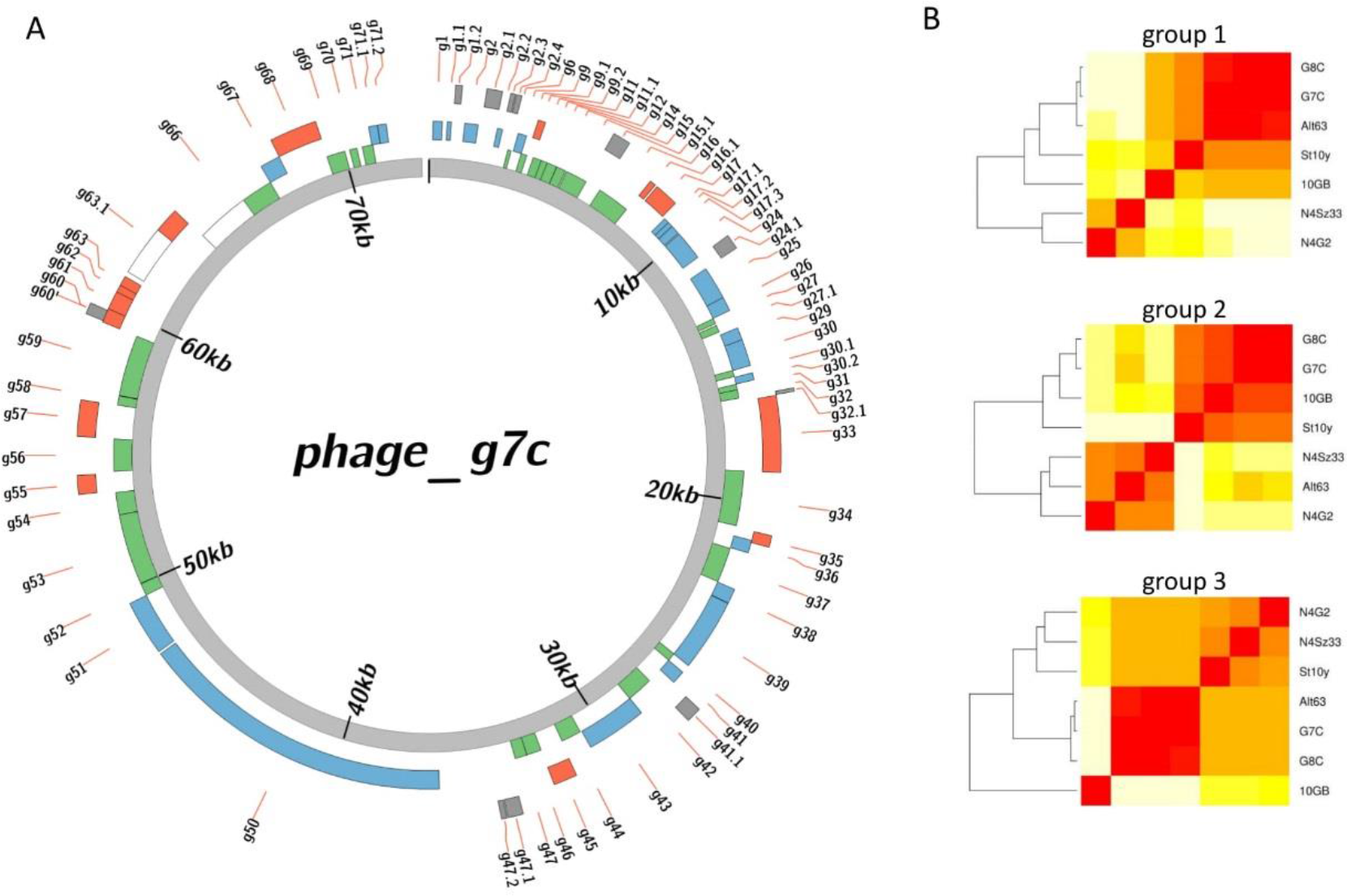
**A.** The distribution of the genes belonging to three identified groups on phage G7C genomic map. Blue – group2, green- group1, red- group-3, grey – the genes not included to local core genome. Only N-terminal parts of the tailspike proteins genes 63.1 and 66 are considered as a part of the local core genome. **B.** Phylogeny of the G7C-set phages based on the subsets of the local core-genome genes belonging to each of the three groups.

To further analyse the properties of each of three clusters we calculated the phylogenies of the phages based on the sequences of the genes of each cluster (Fig. 5B). One can see that the phylogeny based on the largest group #1 fits well to the whole genome phylogeny calculated using Jspecies algorithm (Fig. 1B). At the same time in phylogenies based on the groups #2 and #3 the positions of the phages St10y and 10GB in respect to the other genomes is altered while the clusterisation of the remaining 5 genomes is similar to the group #1 – based phylogeny (Fig. 5B). This is indicative for mosaic nature of the genomes of St10y and 10GB phages both isolated in 2010. To be able to visualize this mosaicism more clearly, we built a heatmap of relatedness of each of 67 local core-genome genes in each of the phages to G7C phage that was taken as a reference (Fig. 6).

**Fig. 6.**
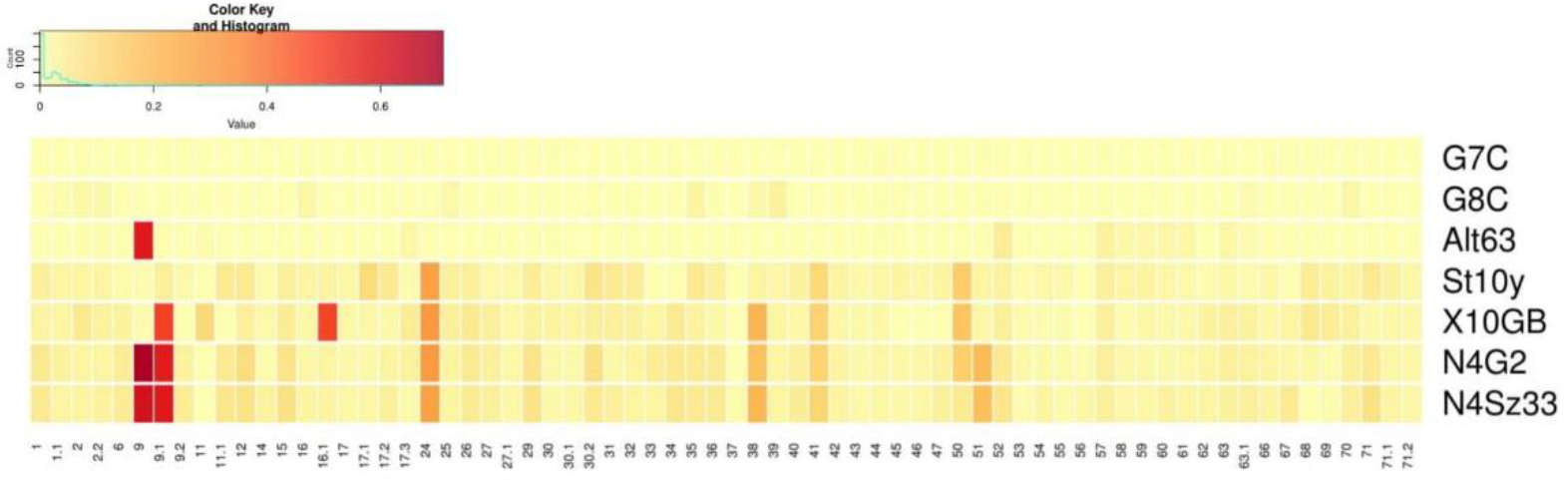
The heatmap of the distances of the genes belonging to the local core genome in each of G7C-set phage from the corresponding genes of the phage G7C taken as a reference.

On this diagram one can see that the distances of the core genome genes from the reference are much stronger correlated between N4G2 and Sz33 or between G8С and Alt63 than between St10y and 10GB. Suggesting that the later two genomes are formed by recent recombination that brought together the fragments of highly related but not identical phage genomes. Interestingly the pattern observed in genomic fragment from gene 52 to gene 63 is similar for the phages St10y, 10GB, N4G2, Sz33 and Alt63 while the rest of Alt63 genome is highly related to G7C reference.

To further, confirm this the local hidden mosaicism we analyzed the distribution of SNP densities along the alignment of our phage genomes against phage G7C using methods developed in (Dixit et al. 2015). The results (Figs. S1-3) suggest that the two phages (Alt63 and G8C) which are the most closely related to the reference strain G7C have around 92% of their genomic regions vertically inherited from their last common ancestor with the phage G7C. These long (up to 33kbp) vertically inherited regions are interspersed with relatively short recombined regions (average length of around 0.6kbp). Comparison of four other genomes (10GB, N4G2, N4Sz33, and St10y) to the reference strain (phage G7C) revealed much shorter uninterrupted vertically inherited genomic regions (the average length of only around 0.5kb). Hence, in the course of the evolution of these phages starting with their common ancestor with the phage G7C, the bulk of their genomic sequences were covered by a mosaic of overlapping recombined segments acquired from multiple donors.

### Estimate of the divergence time of G7C-set bacteriophage genomes

The times of divergence of six selected genes were estimated using BEAST algorithm (Drummond and Rambaut 2007). For all of the genes except gene 39 the oldest bifurcation age was estimated within 11 – 25 years prior the last sampling in 2013(Figs. S4-9). For gene 39 the separation of the phage 10GB allele from the other phages was estimated to occur 43.5 years before the last sampling.

### Genetic alterations of phage G7C genome during in vitro coevolution with E. coli 4s in the laboratory microcosms

In order to compare the genetic alterations of the virus genome that occur during the co-evolution of G7C phage and its host in vivo and in the simplified system, we conducted two experiments. In one experiment, the phage was let to co-evolve with the host in the liquid periodical culture for 7 and 12 passages (ca. 50 - 100 host generations) and then the viral metagenomes were sequenced. Surprisingly no SNPs present in more than 10% of the sequence reads were found.

In the other experiment we used the co-evolution on solid medium that was supposed to provide some spatial complexity that may influence the evolution rates and trajectories (Morgan et al. 2005; Kerr et al. 2006; Vos et al. 2009; Roychoudhury et al. 2013; Ashby et al. 2014; Hesse et al. 2015; Möbius et al. 2015). The material from 60 G7C plaques grown on the E. coli 4s lawn was transferred to the LB plate and the growth of the cultures (metastable phage-bacteria associations) was observed. In course of sequential passages the number of associations that continued to produce lysis zones on the E.coli 4s lawn declined (Fig. S10) and after the passage #11 all of them ceased to produce the detectable phage. However, 4 cultures continued to produce the phage detectable on E. coli 4sAs lawn. These (meta) stable associations were maintained for more than 20 passages and continued to produce the phage. These phages were able to grow on E. coli 4sAs mutant strain but were all noninfectious for the parental wild type (wt) strain. We selected several such isolates and compared their adsorption to E. coli 4s and E.coli 4sAs. No adsorption could be seen on wt for any of the virus isolates while 4sAs strain cells adsorbed all the phage strains rapidly. To identify the genetic basis for this phenotype we sequenced the genome of one of the E. coli 4sAs specific phage isolate named ph8810. Only one SNP leading to the single aa substitution R421P in gp63.1 tail spike was observed (Fig.). We analysed gene 63.1 sequences in 4 more phage isolates obtained from our associations that were indistinguishable phenotypically from ph8810. Several other aa substitutions were observed (data not shown) but only R421P was conserved in all the lineages.

As it was expected the only detected source of the diversity during phage adaptation in course of in vitro co-evolution with its host were spontaneous point mutations.

## Discussion

### The G7-like phages

The *E. coli* phage N4, the prototype species of the group of N4-related viruses was a database orphan for many years (Kazmierczak et al. 2002). However, over the last decade, numerous N4-like viruses that infect Gram-negative bacteria were isolated and sequenced (Faidiuk et al. 2017; Golomidova et al. 2007; Cai et al. 2015; Patel et al. 2015; Li et al. 2016; Ma et al. 2016; Shigehisa et al. 2016; Bhattacharjee et al. 2017; Katharios et al. 2017; Buttimer et al. 2018), and for reviews see (Chan et al. 2014; Wittmann et al. 2015). The N4-related sequences are also ubiquitous in human fecal metagenomes and viromes (Waller et al. 2014). *E. coli* G7C-like phages were previously identified as indigenous components of the horse intestinal viral community (Golomidova et al. 2007; Kulikov et al. 2012) and this work.

### The horse as a natural experimental system

In order to characterize this virus group further it is useful to observe it’s evolutionary trajectory. Ideally this would best be carried out in situ and the physiology of horse digestive tract makes it an attractive model system to study *in situ* bacteriophage ecology adaptation. This is because the microbial digestion of the cellulose-containing food takes part in the lumen of the horse hindgut where the contents are well mixed by the peristaltic movements. The large volume of the gut also limits the impact of the mucosa associated bacteria have over the overall community structure. Moreover, the horse diet is very stable and the average time between food intake (and between defecation) is much shorter than the 48-72 h food retention time in the gut (Hintz and Cymbaluk 1994). The horse gut therefore functions as a natural chemostate populated by a complex high-density community that includes bacteria, archaea, protozoa and associated viruses. *E. coli* is a minor but permanently present component of the horse gut ecosystem. In our previous work we demonstrated that *E. coli* diversity at the strain level in the horse feces is extremely high, reaching several hundreds of lineages per sample (Golomidova et al. 2007; Isaeva et al. 2010). At the same time, all the coliphages cultured by us from *E. coli* feces to date appear to be virulent (Golomidova et al. 2007; Kulikov et al. 2014).

### G7C-set represents an endemic viral population

We studied this G7C-like phage group and the external environment(s) surrounding them, which presumably act as opportunities to connect the individual microbiomes and viromes. The whole-genome based phylogeny of G7C-set indicates that these phages are much closer related to each other than to their closest homolog found in GenBank – PhAPEC7 – another G7C like virus. Other features that, the majority of our viruses encoded that make them distinct from know viruses are; 4 out of 7 contain a G7C-type of gp63.1, a tailspike with a rare deacetylase activity (Prokhorov et al. 2017) – this feature has never found in any N4-related phages outside of this set (the closest BlastP hits for G7C gp63.1 are for *Morganella* and *Enterobacter* prophage proteins). These results strongly support the hypothesis that the divergence observed between our isolates originated from local microevolutionary processes that take place in this particular horse population and in contrast to the alternative hypothesis that this diversity was elsewhere and collected due to the settlement of the externally acquired phages in the viromes of the animals studied.

Further evidence for the endemic nature of this G7C-set is that all phages have very narrow host ranges – and only infect the *E. coli* 4s strain and its derivatives, and that no N4-like phages from other horse populations could be isolated to grow on this strain.

### Estimation of the age of the viral cluster

The G7C set forms a tight cluster on a whole genome based phylogentic tree (Fig. 1B) and appears to be endemic to this particular horse population. The timescale over which the evolutionary process have occurred is therefore likely to be over the last 25 years – which is when the horse yard was established, or possibly more recently, if they were brought in by a newly acquired horse brining the ancestral phage group in. We do know however that a representative was seen in 2006. This timeframe is further supported by the BEAST analysis, which reconstructs phylogenies and estimates the age of the branches in order to obtain a molecular clock. This analysis gives an estimated divergence time for most of the markers analysed of well within 25 year range prior to the last sampling (2013). The only exception to this was the g39 based BEAST where the branch that separates the 10Gb from the rest of the G7C set is estimated to be 43 years. However, in this phage gene 39 carries a non-self-splicing insertion. The selective force caused by adaptation to this re-arrangement (i.e. putative transition from monomeric to two subunits protein) could increase the SNP density in this gene increasing the apparent separation time. The estimated times of separation of gene 39 sequences from the other phages are all below 17 years (Fig. S9).

### Lateral gene transfer in the G7C-set

While the whole-genome based phylogeny of the G7C-set isolates generally follows the isolation time (Fig. 1), the pattern of the detected recombination events (Table 1) does not follow this phylogeny (Fig. 2). This indicates that many of the observed large polymorphisms (modular replacements, deletions or insertions of large sequence fragments) emerged relatively recently. This fits well the hypothesis that all the genotypes of G7C set originated in situ, and were not acquired from the external sources. It also highlights the importance of modular mechanisms in driving diversity during the evolution of the G7C-set. In fact, in average more than half of the divergent nucleotide position in pairwise alignments G7C-set phage genomes are located within the loci affected by the lateral gene transfer of unrelated sequences. This makes a striking contrast to the genome dynamics observed in experimental macrocosms populated by simplified microbial communities (Akusobi et al. 2018; Buckling and Brockhurst 2012; Golais et al. 2013; Petrie et al. 2018; Scanlan et al. 2011) and this work; see also Supplementary Discussion.

The local core genome of this phage population is also highly mosaic due to frequent recombination events. However these recombinations largely take place between closely related viral genomes. This finding is in agreement with recent work (Gregory et al. 2016; Marston and Martiny 2016) demonstrated that in marine T4-related cyanophages the on-going recombination between highly similar (> 93% identity) viral genomes may be one of the major source of diversity.

The invasion of the introns or intron-like non spliced sequences observed in two different essential core genes suggest that in the some contexts (i.e. under the influence of specific genetic pool of a given microbial community) the core genes may undergo significant changes. This may contribute to core genome erosion that is easily observable at much longer evolutionary distances (Mann et al. 2005; Cazares et al. 2014). Interestingly the insertion of a mobile non-sliceable sequence into the gene 39 of phage 10GB led to separation of the corresponding protein, the DNA-polymerase into two distinct polypeptides that appear to act as separate subunits. Clearly further experimentation is needed to confirm that both these polypeptides are necessary for enzyme activity. However, the rapid change of the tertiary structure of an essential gene resulting from the activity of a mobile element is an interesting example of a mechanism driving the erosion of the core genome of a phage group.

## Conclusions

To summarize our data, we can conclude that one of the major factors shaping the evolutionary trajectories during short-term evolution of G7C-related phages in horse – associated microbial system is the access to the large genetic pool of the complex natural microbial community. This differs from the process of evolutionary adaptation of the phage genomes *in situ* from what can be seen or even expected in the simplified laboratory settings.

At the same time, we need to highlight that observation that the core viral genome in this phage-population does not evolve only vertically but is also subject to frequent recombination events. However, these recombination events are largely limited to recombination between closely related genotypes, most likely due to stronger selection pressure.

This data provides interesting new examples to support the reconceptualization of the bacteriophage population core-genome. This is ultimately likely to lead to a deeper understanding of bacteriophage phylogeny. Moreover, the consideration of the local population core metagenomes (and not the individual genomic sequences) as the evolving entities can be justified. Therefore, the development of the approaches to use the meta-sequences of the local phage population core genomes as a short time-scale molecular clock can be required. Additional work is necessary to build a more detailed picture of the phage adaptation processes resolved in time. This may be achieved using the combination of the strategy based on phage isolation applied in this study that allows to examine experimentally the physiological consequences of the alterations observed, and of novel metagenomic approaches using single molecule sequencing techniques that will allow to completely asses the phage genotypes diversity at each time point.

## Materials and methods

### Bacterial strains and phages

This study exploited the environmental isolate *E. coli* 4s and its derivative 4sI lacking the o-antigen O-acetylation, these have been characterized previously (Knirel et al. 2015).

### Sampling and bacteriophage isolation

The samples of horse feces were collected at Children equestrian sportive center in Neskuchny garden in Moscow. The sampling and bacteriophage isolation from the horse feces was performed as it described earlier (Golomidova et al. 2007). The first sampling took place in 2006 in course of previous study Golomidova et al. 2007) The bacteriophages G7C and G8C were isolated by a chance as representative phages for E. coli 4s strain. Phage Alt63 was then discovered as a contaminant of the original G7C stock (Kulikov et al. 2012). The samples obtained in 2010 and 2013 were processed in similar way. The phage plaques obtained were screened using PCR for conserved g50 sequence to detect G7C-related phages. The positive phages where plaque-purified, the lysates were grown, the DNA was extracted and EcoRV restriction profiles were determined. The phages with any detectable differences in restriction profile were sequenced.

### PCR and primers

Primers used in this work are listed in the Table S1. The PCR reactions were performed by standard protocols using Taq (Chomczynski and Sacchi 1987) and Pfu polymerases (the detail of the PCR protocols are available from the authors). For reverse – transcription PCR the RNA was extracted from the infected cells as described by (Chomczynski and Sacchi 1987), cDNA was synthetized using GoScipt Reverse transcriptase (Promega, USA) according to the manufacturer’s instructions with the primers same primers that were then used for the PCR. Gene 16 from phages N4G2, Sz33, and gene 39 from phage 10Gb were cloned into pGEM-T vector according to the manufacturer’s recommendations. Then the clones with the appropriate orientation of the insertion for expression under the control of t7 promoter were selected.

Gene 63.1 from bacteriophage Alt63 was cloned into pET23a using the restriction sites XbaI and XhoI to produce the expression vector p23rA63.1HT.

### Bacteriophage DNA isolation and sequencing

Bacteriophages were purified by repeated single plaque isolation and DNA was prepared as described previously (Kulikov et al. 2014). For library preparation, amplified cDNAs (100 ng of each sample) were fragmented by 200-300 bp using the Covaris S220 System (Covaris, Woburn, Massachusetts, US). Next, the Ion Xpress Plus Fragment Library Kit (Life Technologies) was employed for barcode shotgun-library sample preparation. To conduct emulsion PCR, the Ion PGM Template OT2 200 Kit (Life Technologies) was utilised. Sequencing was performed in accordance with manufacturer protocol for the genome analyser Ion Torrent PGM (Life Technologies), using an Ion 318 chip and Ion PGM Sequencing 200 Kit v2 (Life Technologies). For de novo phages genome assembler, Newbler (Roche Diagnostics, Basel, Switzerland) was used in default mode.

### Bacteriophage phylogeny reconstruction

To reconstruct the whole genome based phylogeny we used an alignment independent algorithm JSpecies (Richter et al. 2016). As an out-group for this phylogenie, we used a complete phage genome of bacteriophage that was the best match found within the NCBI blastN search using phage G7C as a query (Fig. 1B).

The estimates of divergence time were done based on BEAST analysis of the selected genes each of which was found in all the phages studied.

To estimate the divergence time of G7C-set bacteriophage genomes phylogenies of for 6 selected core genes were calculated using a Bayesian likelihood-based algorithm implemented in beast v.1.8.4 (Drummond and Rambaut 2007) Genes 33 (rIIA-like protein), 39 (DNA polymerase), 59 (major capsid protein), 68 (structural protein) and conserved N-terminal moieties of receptor recognition proteins genes 63.1 and 66 were used. The SRD06 codon-based substitution model (Shapiro et al. 2006) was used with a random local clock. A coalescent constant size tree prior was used because in preliminary runs it yielded a higher effective sample size (ESS) of the posterior probability values than coalescent: exponential growth or coalescent: expansion growth tree priors. Each analysis was run for 100 million generations to achieve an ESS >200. The trees were sampled every 100 generations. Trees were annotated with TreeAnnotator v.1.8.4 with a burn-in of 10% and visualized with FigTree v.1.4.3 (http://tree.bio.ed.ac.uk/software/figtree/). Tree roots were calculated by beast software, and no outgroup was used for rooting.

### Analysis of the ‘local core genome’

The genes or part of the genes that are present in all seven genomes of G7C-set phages were termed 'the local core-genome'. All core genes were aligned with muscle. The distance between genes was calculated in R with dna.dist function from package ape (K80 evolutionary mode). Then spearman correlation of distances was calculated resulted in a matrix of correlation coefficients between different genes. This matrix was clustered with hclust function with ward.D agglomeration method and obtained tree was divided into four clusters with cutree function.

### In vitro co-evolution experiments

Co-evolution of the phage with the host in the liquid macrocosmes 10 ml of LB medium was inoculated with *E. coli* 4s culture and grown to the OD_600_ = 0.6. The culture was infected with G7C phage at the multiplicity of infection of 10^−3^ and incubated overnight at 37°C with agitation. Next day culture was diluted 100 times and the grown overnight in the same conditions. The whole experiment was set in triplicate. In one of the microcosms the phage was lost after 5 passages that was confirmed by PCR for gene 50 sequence. The phage fraction was isolated from the remaining two parallels at the passages 7 and 12, the viral DNA was extracted and sequenced as the metagenomes.

The SNP detection was performed in the metagenomes sequences compared to the original phage stock dataset. We set up 10% SNP coverage threshold level to avoid the misinterpretation of the sequence errors as biologically meaningful information.

Co-evolution of the phage with the host on the solid medium was followed in the experimental set up described previously (Letarov and Krisch 2013). Briefly, the material from 60 phage G7C plaques obtained on *E. coli* 4s lawn was transferred to the fresh LB plate. The growth of bacterial cultures (Pseudolysogenic associations, PA) was observed in all the cases. The cultures were transferred by the toothpicks to the fresh plate and simultaneously on the plate with fresh lawn of *E. coli* 4s. The passages were repeated 20 times. The percentage of the cultures producing detectable phage decreased gradually with the number of the passages and all of them ceased to produce such phage by passage #11. However some of the cultures obtained in this experiment were later noticed to produce plaques when they were used for production of the lawn (autoplaquing). The cultures were then suspended in phage neutralizing solution (ref) diluted and plated to obtain the single colonies. Screening these colonies for the ability to support the growth of the phage present in the autoplaquing cultures we identified a derivative of the parental *E.coli* 4s strain named *E. coli* 4sAs.

In this work the experiment with co-evolution similar to the one described above was then repeated and *E. coli* 4sAs lawns were used in addition to *E. coli* 4s for control of the phage production by the PA.

## Acknowledgements

The work was supported by RFBR grant 18-29-13029 (bioinformatical work) and RSF grant 15-15-00134P (the expression and the analysis of the activity of gp63.1 protein from Alt63 phage).

