## Supplementary Information for "Phages associated with horses provide new insights into the dominance of lateral gene transfer in virulent bacteriophages evolution in natural systems"

**Analysis of vertically inherited (clonal) and recombined segments in phage genomes**

We carried out the analysis of vertically inherited (clonal) and recombined regions in phage genomes using methods developed in (Dixit et al., 2015) for different strains of E. coli. Genomes of six phage strains (10GB, Alt63, G8C, N4G2, N4Sz33, and St10y) have been aligned to the reference strain G7C as described before. The local sequence divergence between each of these strains and G7C has been quantified by the number of SNPs within consecutive 100bp genomic segments. Because of differences in genome lengths of individual strains the number of such segments slightly varies from strain to strain and ranges between 694 (69.4kbp total length) and 717 (71.7kbp total length).

Figure S1 shows the histogram of the number of SNPs (per 100bp) separating different strains from the reference strain G7C. The genomes of strains Alt63 and G8C (but not those of the remaining four strains) when aligned against the genome of the strain G7C show a clear separation between vertically inherited segments with 0 SNPs (a sharp peak at the left part of the distribution) and the exponential distribution of the number of SNPs in the rest of the genome. Similar form of the distribution has been reported in (Dixit et al., 2015) for genomes of recombining bacterial (E. coli) strains. This form of the distribution allows one to separate the vertically inherited (clonal) genomic segments in the “clonal peak” (which all have exactly zero SNPs in our case) from the rest of the genome composed of a variable number of recombined segments.

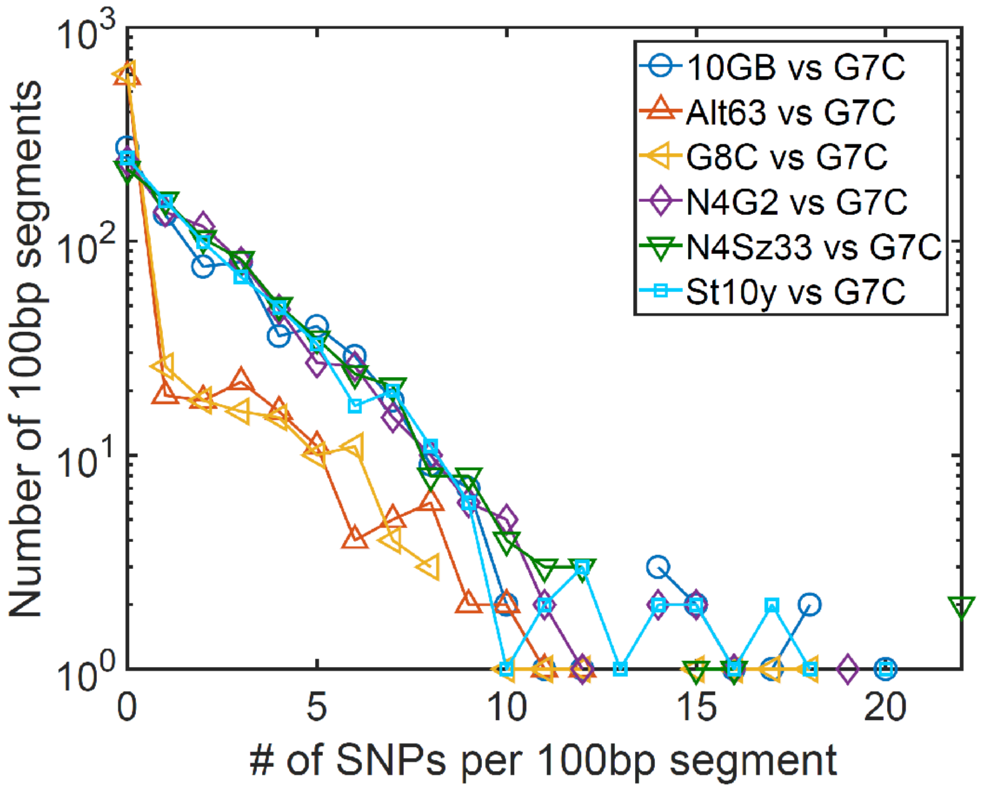


Figure S1. The histogram of the number of SNPs (per 100bp genomic segment) separating each of the six strains (10GB, Alt63, G8C, N4G2, N4Sz33, and St10y) from the reference strain G7C. Strains Alt63 and G8C (but not the other four strains) show a clear separation between vertically inherited segments with exactly zero SNPs (peak at the left) and recombined segments with 1 or more SNPs. The number of SNPs in such recombined segments approximately follows an exponential distribution. The histogram is qualitatively similar to that reported for recombining strains of E. coli (see Fig.2 in (Dixit et al., 2015)).

To better separate the vertically inherited (clonal) and recombined segments we considered their positions along the chromosome. As established in (Dixit et al., 2015), the SNP density within recombined regions is highly inhomogeneous. Hence, such regions might contain isolated segments with low levels of divergence. To prevent erroneously labeling such segments as "vertically inherited (clonal)" we implemented the following rule: an isolated 100bp segment with zero SNPs surrounded on both left and right by 100bp segments with 1 or more SNPs is reclassified as a recombined segment. We identified and corrected 7 (and 4) such segments in Alt63 vs G7C (and G8C vs G7C) alignments respectively.

On the other hand, some of the clonal segments contain isolated de novo mutations that have originated since the pair of strains diverged from their last clonal ancestor. To avoid erroneously labelling such mutated vertically inherited regions as recombined we apply the following rule. An isolated 100bp segment with exactly 1 SNP surrounded on both left and right by 100bp segments with 0 SNPs is reclassified as a clonal segment. The number of reclassified segments of this type in different genome comparisons ranged between 1 and 4.

Clonal segments form the majority of all 100bp segments in Alt63 vs G7C and G8C vs G7C alignments with 581 out of 694 segments (83.7%) and 605 out of 715 segments (84.6%) respectively. The remaining segments have been recombined in at least one of the two aligned genomes. Assuming equal recombination rates in each of two genomes, we estimate that around 92%=84%+16%/2 of each of the three genomes (Alt63, G7C and G8C) has been vertically inherited from their last clonal ancestor.

We analyzed the length distribution of uninterrupted regions of consecutive clonal segments on the chromosome (in what follows referred to as "clonal regions"). Our interpretation of such regions as vertically inherited was corroborated by the observation that for Alt63 and G8C compared to G7C, uninterrupted clonal regions tend to be long. The longest clonal regions are 32.9kbp and 24kbp and the average length of clonal regions is 3.4kbp and 2.9kbp for Alt63 vs G7 and G8C vs G7C comparisons respectively. The longest clonal regions in the other four phage genomes were significantly shorter and ranged between 1.7kbp and 2.9kbp. The average length of clonal regions in each of the four phage strains varied between 0.48kbp and 0.58kbp. This is consistent with a model in which the entire chromosome initially forms one "clonal region" gradually covered by recombined segments resulting in a mosaic of clonal and recombined regions. Most recombination events land within a longer clonal regions thereby breaking it up into two parts. This quickly reduces the maximal and average lengths of uninterrupted clonal regions. The cumulative distribution of lengths of uninterrupted clonal regions is shown in Fig. S2.

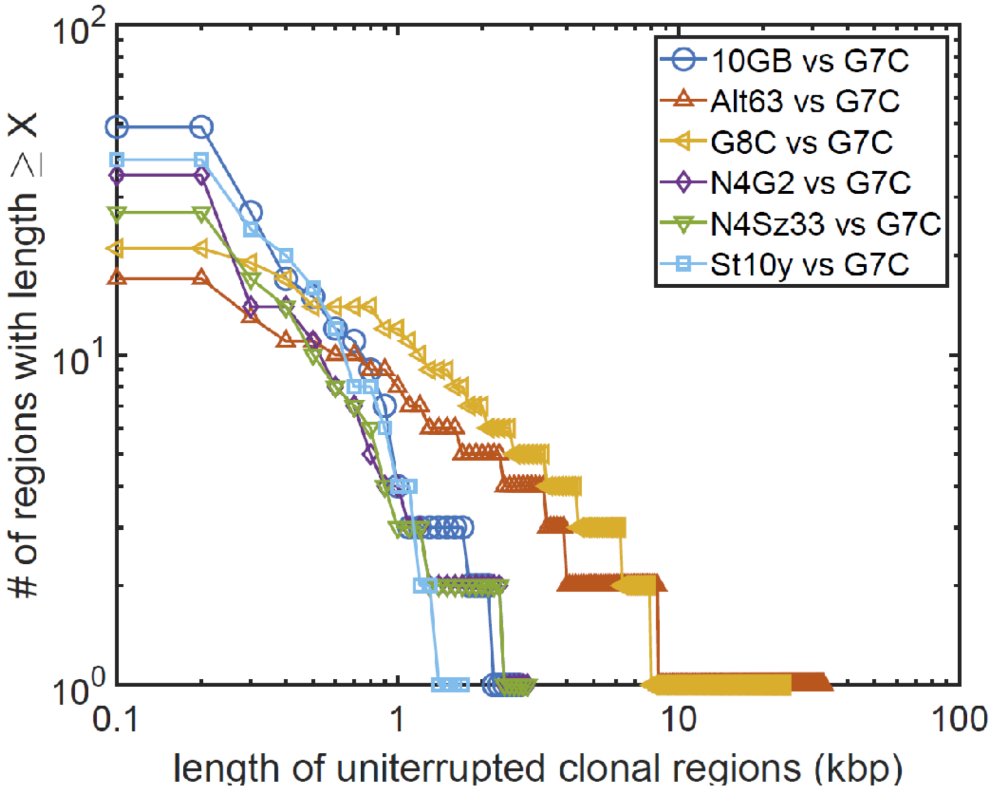


Figure S2. The cumulative distribution of lengths of uninterrupted clonal regions in comparisons of the six strains (10GB, Alt63, G8C, N4G2, N4Sz33, and St10y) with the reference strain G7C. Strains Alt63 and G8C have dramatically longer uninterrupted clonal segments up to 32.9kbp. Note the logarithmic scale on both axes.

We then analyzed the length distribution of uninterrupted regions of consecutive recombined segments on the chromosome (in what follows referred to as "recombined regions"). These regions are considerably shorter than their clonal counterparts. The longest uninterrupted recombined region in our pairwise comparisons ranges between 1.8kb and 7.1kbp, while the average length of uninterrupted recombined regions ranges between 0.52kbp and 2.1kbp. Contrary to clonal regions, longer average recombined regions were observed in four more evolutionarily diverged genome pairs. This is likely due to overlaps between recombined segments incorporated in separate individual recombination events (hidden mosaicism) extending the length of uninterrupted recombined regions. Two closely related genome pairs (Alt63 vs G7 and G8C vs G7C) have, respectively, 17 and 21 putative recombined segments. Assuming equal recombination rates, about half of those segments have been recombined into each of the two genomes since they diverged from their last clonal ancestor. The donor strains of the observed recombined regions had as much as 18% local genomic sequence divergence from the recipient strains (see the largest SNP densities in Fig. S1). The average length of recombined regions in two closely related pairs of strains is 0.66kbp (Alt63 vs G7) and 0.52kbp (G8C vs G7C). The average length of clonal and recombined segments in all six strains is consistent with a model in which a single recombination event on average inserts 0.64 +/- 0.14 kbp of the donor genome into a random position in the recipient genome. The cumulative distribution of the lengths of uninterrupted recombined regions is shown in Fig. S3.

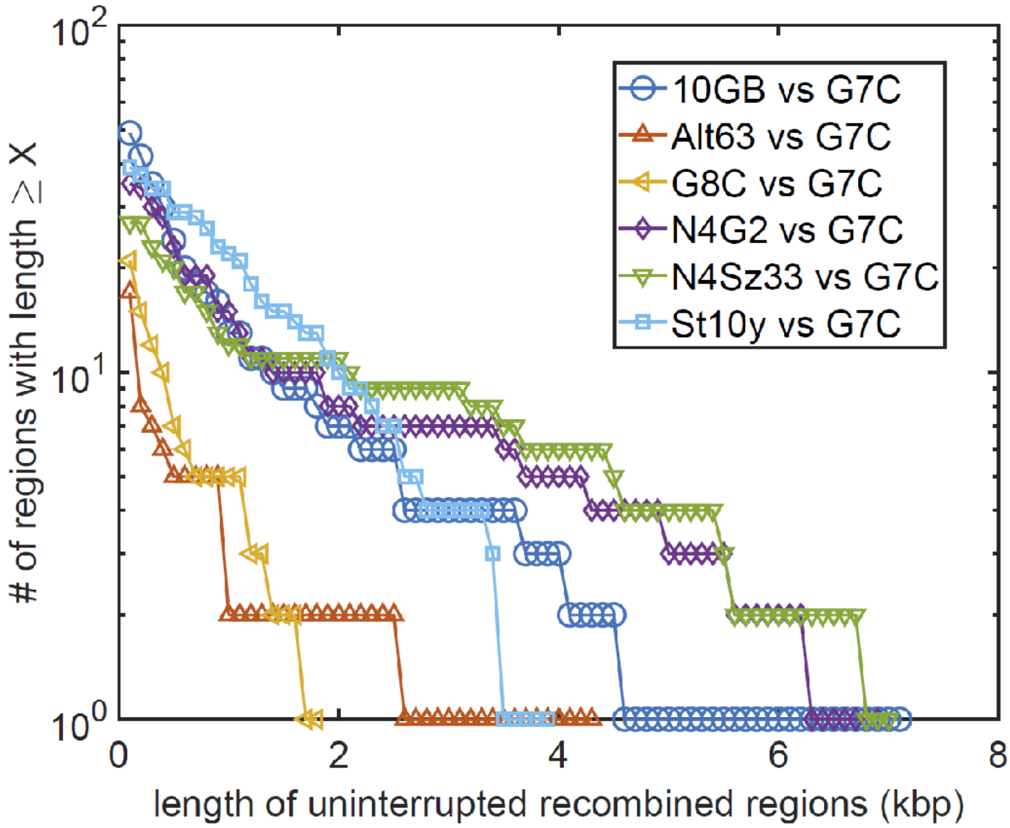

Figure S3. The cumulative distribution of the lengths of uninterrupted recombined regions in pairwise comparisons of the six strains (10GB, Alt63, G8C, N4G2, N4Sz33, and St10y) with the reference strain G7C.
Note that uninterrupted recombined regions are dramatically shorter than their clonal counterparts shown in Fig. S2. Furthermore, uninterrupted recombined regions on average get longer in more distantly related pairs of strains due to overlaps between two or more independently inserted recombined segments. Note the logarithmic scale of the y-axis and linear scale of the x-axis (as opposed to the logarithmic x-axis used in Fig. S2)

**Supplementary Discussion**

The G7C-set phages as well as other cultured N4-like viruses are strictly virulent (i.e. unable for lysogeny) and have to be maintained in the natural habitats *via* their multiplication in the lytic cycle. The second order kinetics of the phage adsorption imposes the minimal host density required for phage maintenance (Kasman et al., 2002; Letarov et al., 2010; Wiggins and Alexander, 1985). The extreme intraspecies diversity of the horse gut *E. coli* population mentioned above suggests that the overall concentration of *E. coli* cells suitable for the multiplication of such a narrow host range phages as G7C-set isolates should be quite low. This allows us to speculate that the long-term existence of G7C-like phages *in vivo* may be ensured by local foci of multiplication of both host and phage. The phage infected microcolonies or biofilm patches may create such an opportunity. The mode of evolution of the bacteriophages in this case should be closer to phage-host co-evolution regimen than to the evolution of the virus only (Buckling and Brockhurst 2012; Golais et al. 2013).

These speculations motivated us to attempt to model of the hypothetical local growth of the phage and the host creating the metastable associations on the solid medium. The analysis of the PAs composition allowed us to identify a simple way for mutual host and phage adaptation that required only a limited number of point mutations both in phage and bacterial genomes. These observations are in good agreement with the other data obtained in some *in vitro* phage-host co-evolution simulation experiments (Buckling and Brockhurst 2012; Golais et al. 2013; Scanlan et al. 2011; Meyer et al. 2012). However the adaptation strategy employed by G7C-like phages *in vivo* appears to be completely different. In these conditions, the impact of the recombination events on the sequence radiation degree (that can be formalized as a number of nucleotide positions altered between any two genomes) appears to be comparable to the effect of the point mutations accumulation. The relatedness of the sub-gene sized modules exchanged in the adsorption proteins gp63.1 and gp66 to various prophages indicates that the recombination with the host genome may be more important for N4-like phages evolution than the recombination with the related phages.

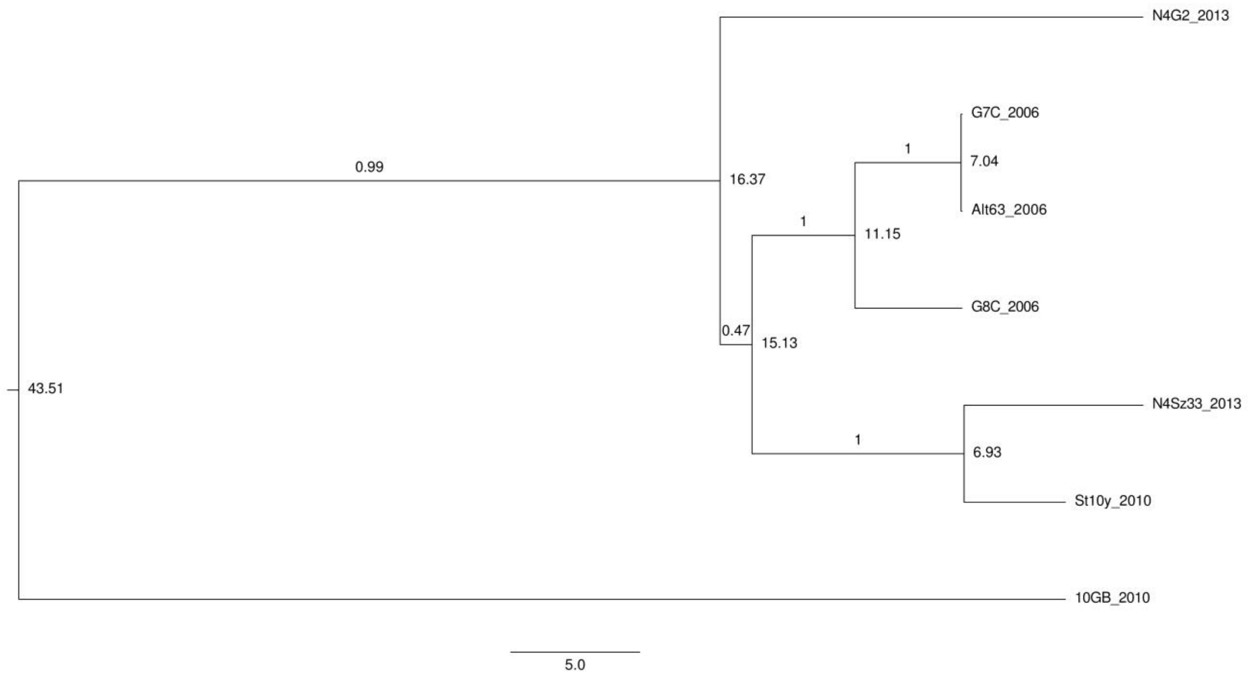


Figure S4. Time-resolved phylogenetic tree for gene 39(DNA polymerase). The numbers in the tree nodes show how long the two lines diverged (years ago, considering that today is 2013 (due to the latest samples)). The numbers in the middle of the branches indicate the probability of relatedness of viruses in the corresponding group. Bar, 5 years.

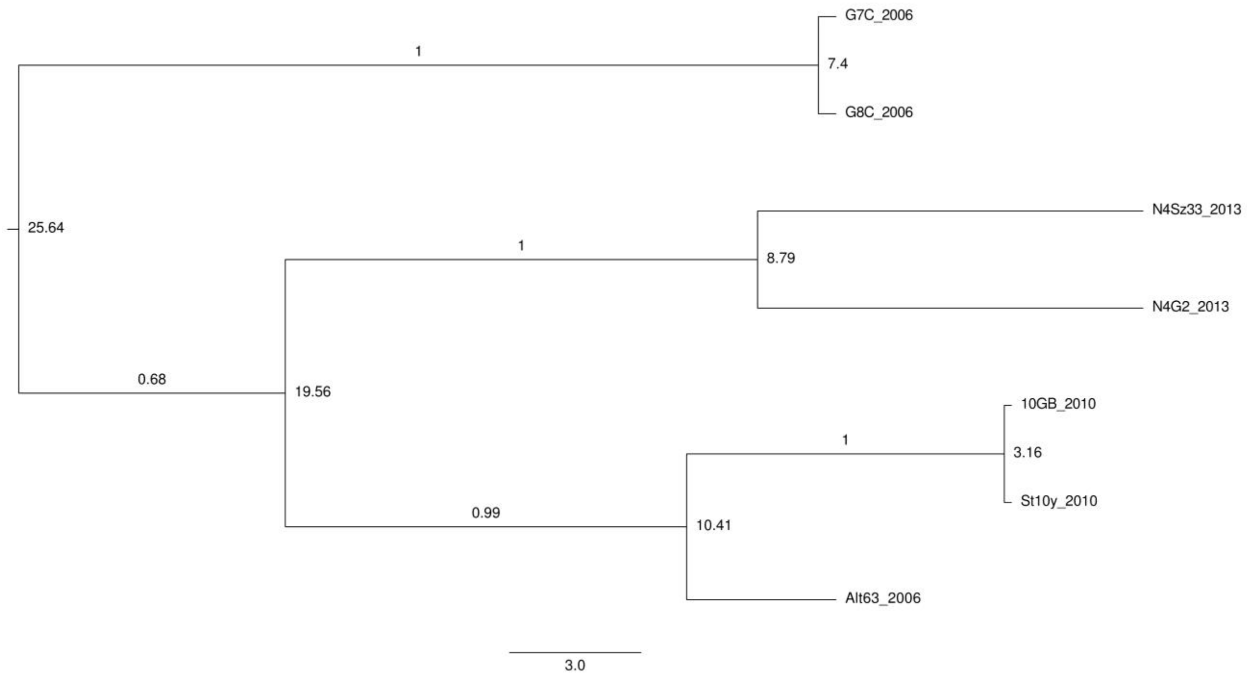


Figure S5. Time-resolved phylogenetic tree for gene 59 (major capsid protein). The numbers in the tree nodes show how long the two lines diverged (years ago, considering that today is 2013 (due to the latest samples)). The numbers in the middle of the branches indicate the probability of relatedness of viruses in the corresponding group. Bar, 3 years.

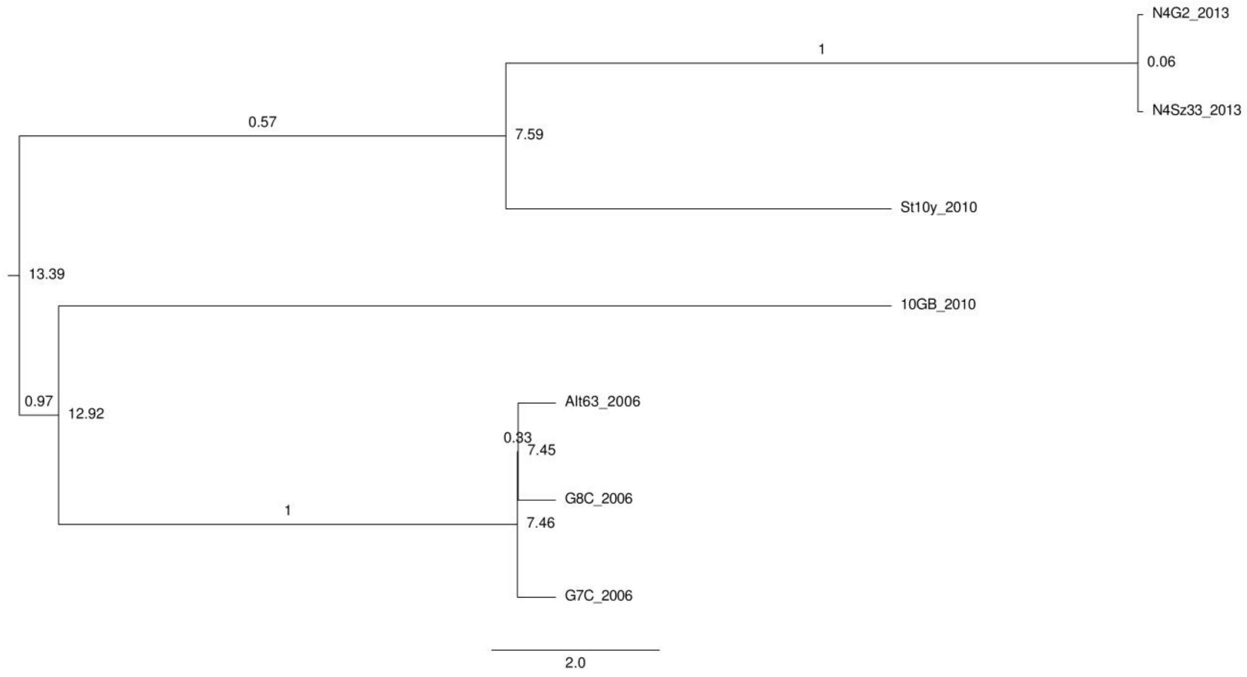


Figure S6. Time-resolved phylogenetic tree for gene 63.1 (conserved N-terminal moieties of receptor recognition protein). The numbers in the tree nodes show how long the two lines diverged (years ago, considering that today is 2013 (due to the latest samples)). The numbers in the middle of the branches indicate the probability of relatedness of viruses in the corresponding group. Bar, 2 years.

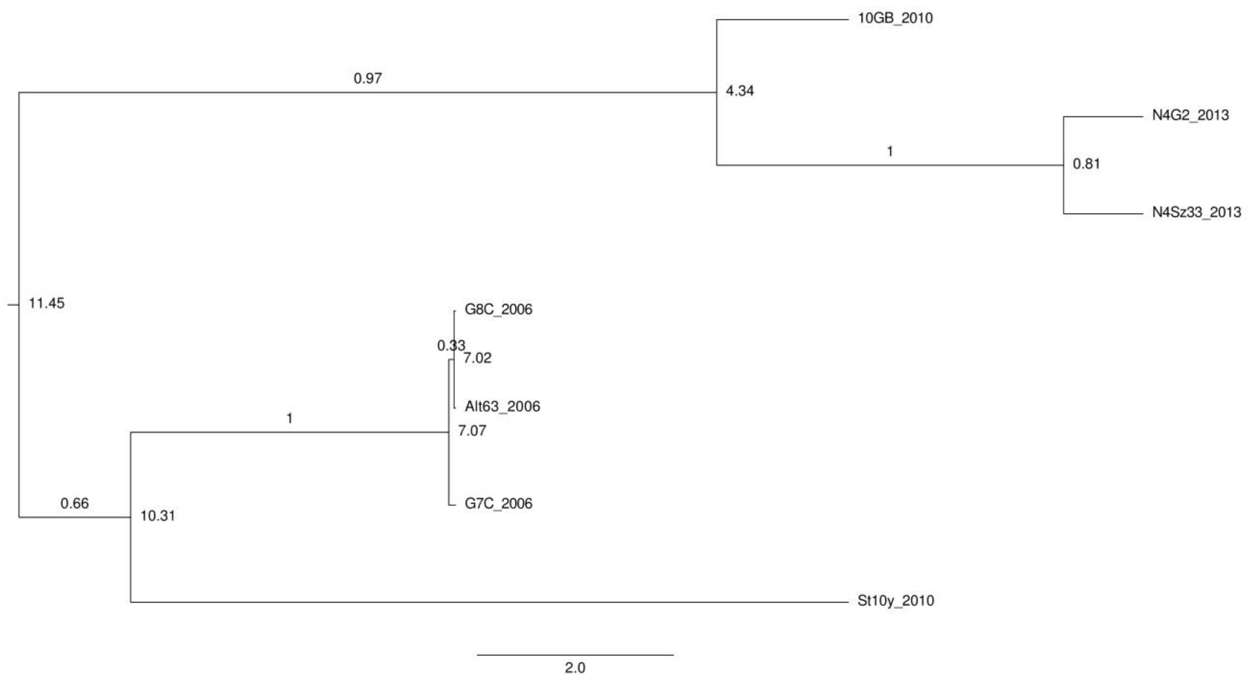


Figure S7. Time-resolved phylogenetic tree for gene 66 (conserved N-terminal moieties of receptor recognition protein). The numbers in the tree nodes show how long the two lines diverged (years ago, considering that today is 2013 (due to the latest samples)). The numbers in the middle of the branches indicate the probability of relatedness of viruses in the corresponding group. Bar, 2 years.

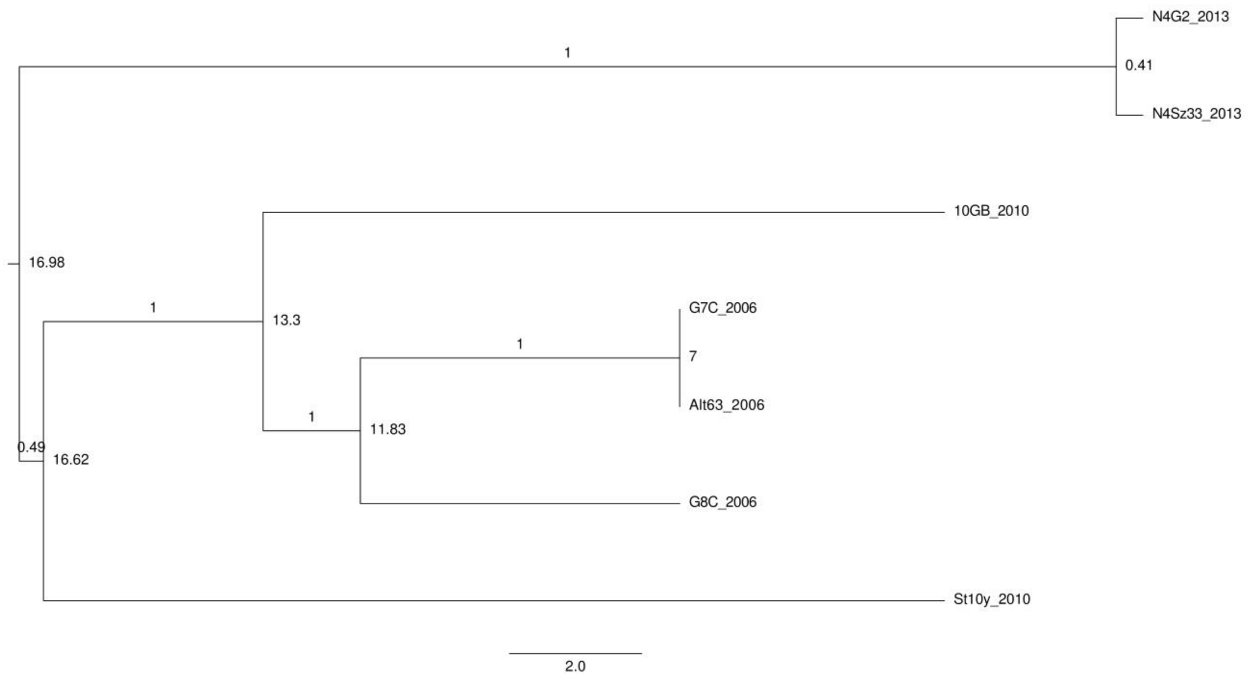


Figure S8. Time-resolved phylogenetic tree for gene 33 (rIIA-like protein). The numbers in the tree nodes show how long the two lines diverged (years ago, considering that today is 2013 (due to the latest samples)). The numbers in the middle of the branches indicate the probability of relatedness of viruses in the corresponding group. Bar, 2 years.

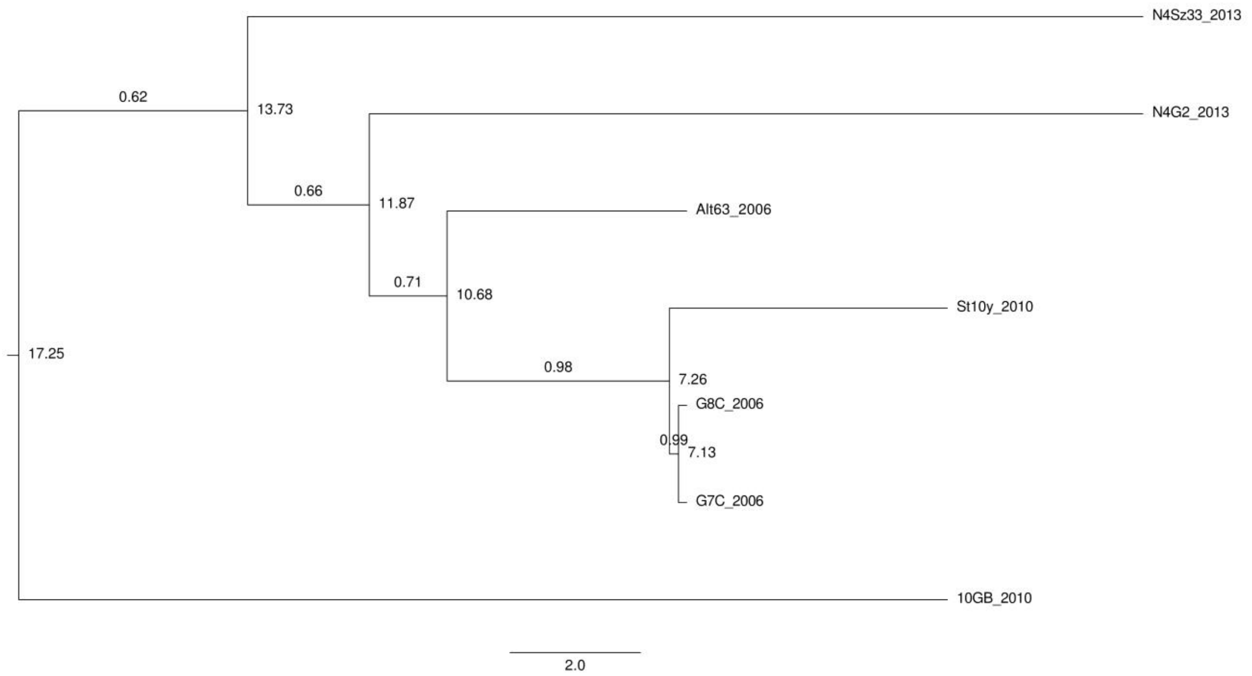


Figure S9. Time-resolved phylogenetic tree for gene 68 (structural protein). The numbers in the tree nodes show how long the two lines diverged (years ago, considering that today is 2013 (due to the latest samples)). The numbers in the middle of the branches indicate the probability of relatedness of viruses in the corresponding group. Bar, 2 years.

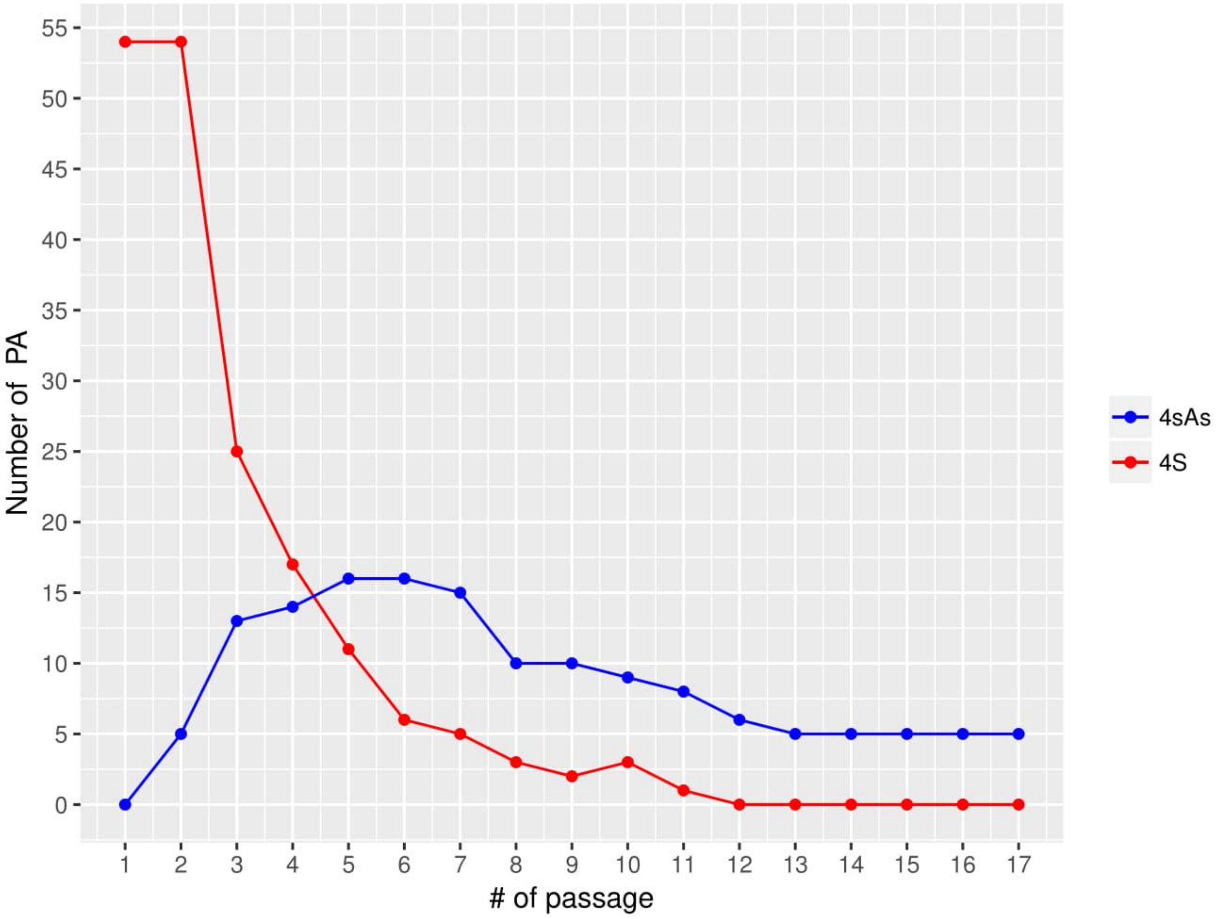


Figure S10.
Phage production by the pseudolysogenic associations (PA) of phage G7C and E. coli 4s host over 17 passages. (Red line) Number of PA producing phage detectable on E. coli 4s lawn. (Blue line) Number of PA producing phage detectable on E. coli 4sAs lawn.

| | Phages | | | | | | Annotation of the affected ORF(s) | Coord mauve by G7C aprox. |
|---|---|---|---|---|---|---|---|---|
| | *G8C* | *Alt63* | *10Gb* | *St10y* | *N4G2* | *Sz33* | | |
| 0 | | | 200 bp insertion before g. 1 | | | | | 118 |
| 1 | | ORF 1.2 deletion | | ORF 1.2 deletion | | | ORF1.2 –hypothetical protein | 780 - 1170 |
| 2 | | | ORF 2.1 deletion | | | | ORF 2.1 putative endonuclease | 1780-2330 |
| 3 | | | ORF2.2 C-term. Extension | | | | ORF 2.2 hypothetical protein | 2170 – 2250 by 10Gb, 2590 – G7C |
| 4 | | | ORF 2.3 deletion, insertion of new ORF 2.4 | | | | ORF 2.3 hypothetical protein | 2600 - 2750 |
| 5 | | | | | | Insertion 350 bp between ORFs 9.1 and 9.2 | ORF 9.1 hypothetical protein, similar to phiEco32 ORF phi32_122 | 3670 |
| 6 | | | | | | insertion of new ORF 14.1 | | 5220 (5120-5360 by Sz33) |
| 7 | | | ORF 15.1 deletion | | | | ORF 15.1 putative homing nuclease | 6090-6670 |
| 8 | | | | | insertion of 800 bp intron with internal ORF into g16 | | gene 16 RNA polymerase 2 subunit A, intron-encoded ORF putative homing nuclease | 7080 |
| 9 | | | 260 bp sequence inserted between ORFs 17.1 and 17.2 | | | | | 9310 |
| 10 | | | ORF 24.1 deletion | | | | ORF 24.1 putative homing nuclease | 10910 - 11400 |
| 11 | | | | | 380 bp sequence inserted between ORF 27.1 and g29 | | | 13720 |
| 12 | | | | ORF 32.1 deletion | | | ORF 32.1 hypothetical protein | 16170 - 16280 |
| 13 | | | | | | fragment of g36 changed | gene 36 conserved hypothetical protein | 21440-21560 |
| 14 | | | insertion of new 500 bp ORF 36.1 | | | | inserted ORF putative homing nuclease, | 21790 |
| 15 | | | insertion of 800 bp sequence with internal ORF into g39 | | | | gene 39 DNA polymerase; internal ORF – putative homing nuclease | 25420 |
| 16 | | | g41 C-truncated, ORF 41.1 deleted. The intergenic spacer remains in its place. | | | | gene 41 – conserved hypothetical protein; ORF 41.1 putative homing nuclease | 26780-27650 |
| 17 | | ORF 47.1 deleted | | | | | ORF 47.1 putative homing nuclease | 33550-34080 |
| 18 | | | internal deletion in the spacer between ORF 47.1 and g50 | | | | | 35100-35270 |
| 19 | | C-terminal part of g63.1 and replaced[a] | | | C-terminal part of g63.1 replaced (similar to Alt63) | | gene 63.1 – enzymatically active tailspike (deacetylase); novel C-terminal domain OPS lyase. | 61200 - 63400 |
| 20 | | | C-terminal part of g66 replaced | | C-terminal part of g66 replaced | | gene 66 – tail spike; novel domains unrelated in three phages | 63850-65-850 |
| 21 | | | insertion of new 350 bp ORF after ORF 71.1 | | | | | 71650 |

Table S1. The list of the modular swapping events observed in the G7C-like phage isolates set. The G7C genome used as a reference.
